# Hypoxia Drives Dihydropyrimidine Dehydrogenase Expression in Macrophages and Confers Chemoresistance in Colorectal Cancer

**DOI:** 10.1101/2020.10.15.341123

**Authors:** Marie Malier, Magali Court, Khaldoun Gharzeddine, Marie-Hélène Laverierre, Sabrina Marsili, Fabienne Thomas, Thomas Decaens, Gael Roth, Arnaud Millet

**Author notes:** Corresponding author: Arnaud Millet MD PhD, Team Mechanobiology, Immunity and Cancer, Institute for Advanced Biosciences, Batiment Jean Roget 3rd floor, Domaine de la Merci 38700 La Tronche, France.

## Abstract

Colon adenocarcinoma is characterized by an infiltration of tumor-associated macrophages (TAMs). TAMs are associated with a chemoresistance to 5-Fluorouracil (5-FU), but the mechanisms involved are still poorly understood. In the present study, we found that macrophages specifically overexpress dihydropyrimidine dehydrogenase (DPD) in hypoxia, leading to a macrophage-induced chemoresistance to 5-FU by inactivation of the drug. Macrophage DPD expression in hypoxia is translationally controlled by the cap-dependent protein synthesis complex eIF4F^Hypoxic^, which includes HIF-2α. We discovered that TAMs constitute the main contributors to DPD activity in human colorectal primary or secondary tumors where cancer cells do not express significant levels of DPD. Together, these findings shed light on the role of TAMs in forming chemoresistance in colorectal cancers and offer the identification of new therapeutic targets. Additionally, we report that, contrary to what is found in humans, macrophages in mice do not express DPD.

## Introduction

Colorectal cancers (CRC) are a leading cause of death worldwide and constitute the third cancer-related cause of death in the United States (Siegel et al., 2020). Chemotherapy is one of the tools used to treat these tumors, however some patients do not respond well to this treatment, resulting in poor prognosis. This chemoresistance is caused by various mechanisms such as drug inactivation, drug efflux from targeted cells and modifications of target cells (Marin et al., 2012; Vasan et al., 2019). Interestingly, the importance of tumor microenvironment in chemoresistance has recently garnered attention. The tumor immune microenvironment (TIME), notably through its innate immune part that is mainly composed of tumor-associated macrophages (TAMs), deserves particular attention (Binnewies et al., 2018; Ruffell and Coussens, 2015). TAMs have been associated with bad prognosis in the case of various solid tumors (Yang et al, 2018) and have been shown to orchestrate a defective immune response to tumors (DeNardo and Ruffell, 2019). It has been suggested that TAMs are reprogrammed by cancerous cells to secondarily become supporting elements of tumor growth (Aras and Zaidi, 2017). The involvement of TAMs in CRC has been controversial, and only recently has their association with a poorer prognosis been recognized (Pinto et al., 2019; Ye et al., 2019). Their implication in chemoresistance, particularly against 5-Fluorouracil (5-FU) a first line chemotherapy in CRC has been reported (Yin et al., 2017; Zhang et al., 2016), suggesting that macrophage targeting could facilitate a way for increasing treatment efficiency. However, the precise mechanisms by which TAMs participate in creating chemoresistance in human colorectal tumors are still unclear. Interestingly, TAMs increase hypoxia in tumor tissues, which is susceptible to favoring chemoresistance in return (Jeong et al., 2019). We also know that hypoxia affects macrophage biology (Court et al., 2017) and could mediate resistance to anticancer treatment and cancer relapse (Henze and Mazzone, 2016). Based on the abundance of macrophages in CRC, we hypothesized that hypoxia could directly modulate macrophage involvement in 5-FU resistance.

## Results

### Macrophages confer a chemoresistance to 5-FU in a low-oxygen environment

In order to evaluate the effect macrophages in the vicinity of the tumor have on chemotherapy, we analyzed the impact on cancer cells growth of conditioned medium (CM) by macrophages containing 5-fluorouracil (5-FU) or none, designed as CM(5-FU) and CM(Ø) respectively. To examine the role of oxygen, we performed these experiments in normoxia (N=18.6% O_2_) and in hypoxia (H=25mmHg ~3% O_2_). Non-conditioned 5-FU strongly inhibited HT-29 and RKO proliferation independently of oxygen concentration (Figures 1A and 1B). CM(Ø) had little effect on the proliferation of colon cancer cells (Figures 1A and 1B). Meanwhile, CM(5-FU 1 μg/mL) inhibited cell growth in normoxia but macrophage conditioning in hypoxia provided complete protection against 5-FU inhibition (Figure 1A, 1B). RKO cells were found sensitive to a lower concentration of 5-FU (0.1 μg/mL). In this state, additionally macrophage conditioning was able to protect against 5-FU not only in hypoxia but also in normoxia (Figure 1B). This observation led us to consider an inactivation mechanism of 5-FU driven by macrophages with increased efficiency in hypoxia. A previously proposed mechanism for macrophage-induced chemoresistance in CRC was related to their ability to secrete interleukin (IL)-6, leading to an inhibition of cancer cell apoptosis (Yin et al., 2017). We first try to verify wether we can confirm the presence of IL-6 in our CM by human monocyte-derived macrophages in normoxia and in hypoxia and found no spontaneous secretion (<10 pg/mL) of IL-6 (data not shown). As the conditioning by macrophages provided a complete protection, we then hypothesized that a direct action of macrophages on 5-FU was the likely mechanism. In order to obtain a molecular explanation of the differing effect under various oxygen concentrations, we performed a proteomic analysis of human macrophages in hypoxia compared to normoxia. Our quantitative proteomic approach revealed that dihydropyrimidine dehydrogenase (DPD) is strongly overexpressed in hypoxia (Figure 1C). DPD (coded by the *DPYD* gene) is the rate-limiting enzyme of the pyrimidine degradation pathway. DPD adds two hydrogen atoms to uracil, with NADPH as an obligatory cofactor, leading to dihydrouracil, which is secondarily degraded to β-alanine under the control of DPYS and UPB1 (Figure 1D). 5-FU is a fluorinated, analogue of uracil reduced by DPD to 5-fluorodihydrouracil in an inactive compound (Figure 1E). We confirmed through immunoblotting the increased expression of DPD in hypoxia when compared to the basal expression in normoxia (Figure 1F). Since the putative action of DPD on 5-FU is related to its enzymatic activity, we also confirmed that DPD expression in hypoxic macrophages was functional, leading to an increase of the dihydrouracil to uracil ratio in the extracellular medium (Figure 1G).

**Figure 1.**
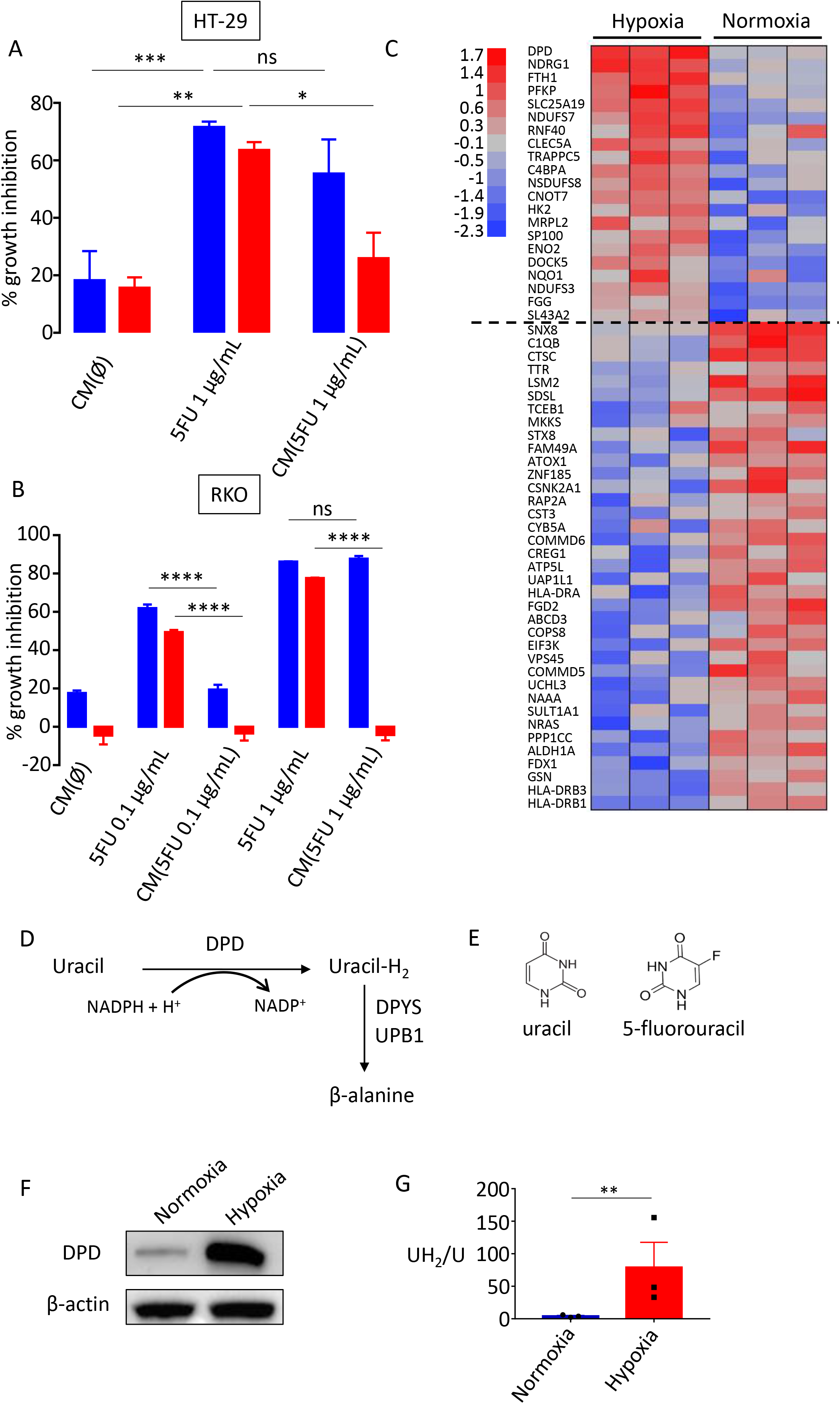
Macrophages confer a chemoresistance to 5-FU in hypoxia. (A) Growth inhibition of HT-29 cells by CM(Ø), 5-FU and CM(5-FU) in normoxia (blue) and in hypoxia (red), 5-FU was used at 1μg/mL (n=3). (B) Growth inhibition of RKO cells by CM(Ø), 5-FU and CM(5-FU) in normoxia (blue) and in hypoxia (red) 5-FU was used at 0.1μg/mL and 1μg/mL (n=3). (C) Protein heat map of macrophages in hypoxia and normoxia. Proteins were selected by a fold change >2 and p-value < 0.01. Proteins were organized according to descending mean z-score of hypoxic proteins. (D) Schematic presentation of the rate-limiting steps of the pyrimidine degradation pathway involving DPD. (E) Chemical structures of uracil and 5-fluorouracil (F) Immunoblot analysis of DPD expression in human macrophages differentiated in normoxia and hypoxia (n= 10). (G) dihydrouracil/uracil ratio measured by HPLC in the macrophage supernatant from macrophages cultured in normoxia and hypoxia (n=3). Error bars represent mean ± sem, *p<0.05, **p<0.01, *** p<0.001, ****p<0.0001, ns=non significant

### Chemoresistance to 5-FU is driven by increased DPD activity in hypoxic macrophages

In order to confirm the inactivation of 5-FU by macrophage DPD and its potential clinical relevance, we analyzed the kinetics of 5-FU degradation in normoxia and hypoxia. As 5-FU is a small molecule, its diffusion in tissue is quite high. Indeed, following four days of oral ingestion of 5-FU the plasmatic concentration was found ~ 1 μg/mL (Zheng and Wang, 2005). The mean 5-FU concentration was 0.411 ± 0.381 (μg/g of the tissue) in the tumor portions of the specimens and 0.180 ± 0.206 (μg/g of the tissue) in the normal portions in colorectal cancers (Tanaka-Nozaki et al., 2001). As TAMs represent 2 −10% of cells in CRC, especially localized in the invasion front (Pinto et al., 2019) and since that 1g of tissue typically contains ~ 10^8^ cells (Wilson et al., 2003), a reasonable estimated ratio in CRC is ~ 10^6^ macrophages/g of tissue. According to 5-FU reported concentration, an estimated physiologically relevant ratio in CRC is 1μg of 5-FU/10^6^ macrophages. We interestingly found that 2 μg of 5-FU could be eliminated by 10^6^ macrophages in hypoxia in less than 24 h and that normoxic macrophages were unable to completely eliminate this quantity in 48 h (Figure 2A). We validated that 5-FU degradation was due to DPD catalytic activity using gimeracil, which is a specific inhibitor of DPD (Figures 2B and 2C). Non-conditioned 5-FU induced death in HT-29 or RKO irrespective of the presence of oxygen and gimeracil, demonstrating no significant DPD activity in these cells (Figures 2D and 2F). We found no significant level of expression of DPD at the protein level, neither in HT-29 nor in RKO cells, irrespective of the oxygen concentration (Figures S1A and S1B). Furthermore, we found a downregulation of the *DPYD* mRNA in HT-29 and in RKO, emphasizing a defective transcription of the DPD gene (Figure S1C). We then confirmed that the pharmacological inhibition of the methyltransferase EZH2 by the specific inhibitor GSK126 led to a detectable level of *DPYD* mRNA in RKO cells (Figure S1D). It has been reported that the transcription factor PU.1 drives the expression of *DPYD* and that EZH2 is responsible for the histone H3K27 trimethylation at the *DPYD* promotor site leading to its downregulation in colon cancer cells (Wu et al., 2016). Whereas CM(5-FU 1μg/mL) induced cell death in HT-29 cells in the case of normoxia, its cytotoxic effect dramatically decreased in hypoxia and this could be reverted by inhibition of DPD activity using gimeracil (Figure 2D). Chemoresistance appeared to be based solely on DPD activity promoted by hypoxia. We observed a similar result using a 3D tumor model growth of HT-29 exposed to CM (Figure 2E), and we confirmed the generality of this mechanism in RKO cells (Figure 2F). In order to confirm that oxygen was the main factor controlling DPD expression in hypoxia, we verified that DPD was not induced or repressed by 5-FU itself (Figure 2G). We also explored wether cancerous cells can modulate DPD expression in macrophages. Using a transwell co-culture between cancerous cells and macrophages revealed no modulation in DPD expression in macrophages (Figure 2H). In order to assess the efficiency of DPD degradation of 5-FU, we also performed a direct co-culture assay between cancerous cells and macrophages and found that macrophages protected cancerous cells from 5-FU in a DPD dependent manner (Figure 2I).

**Figure 2.**
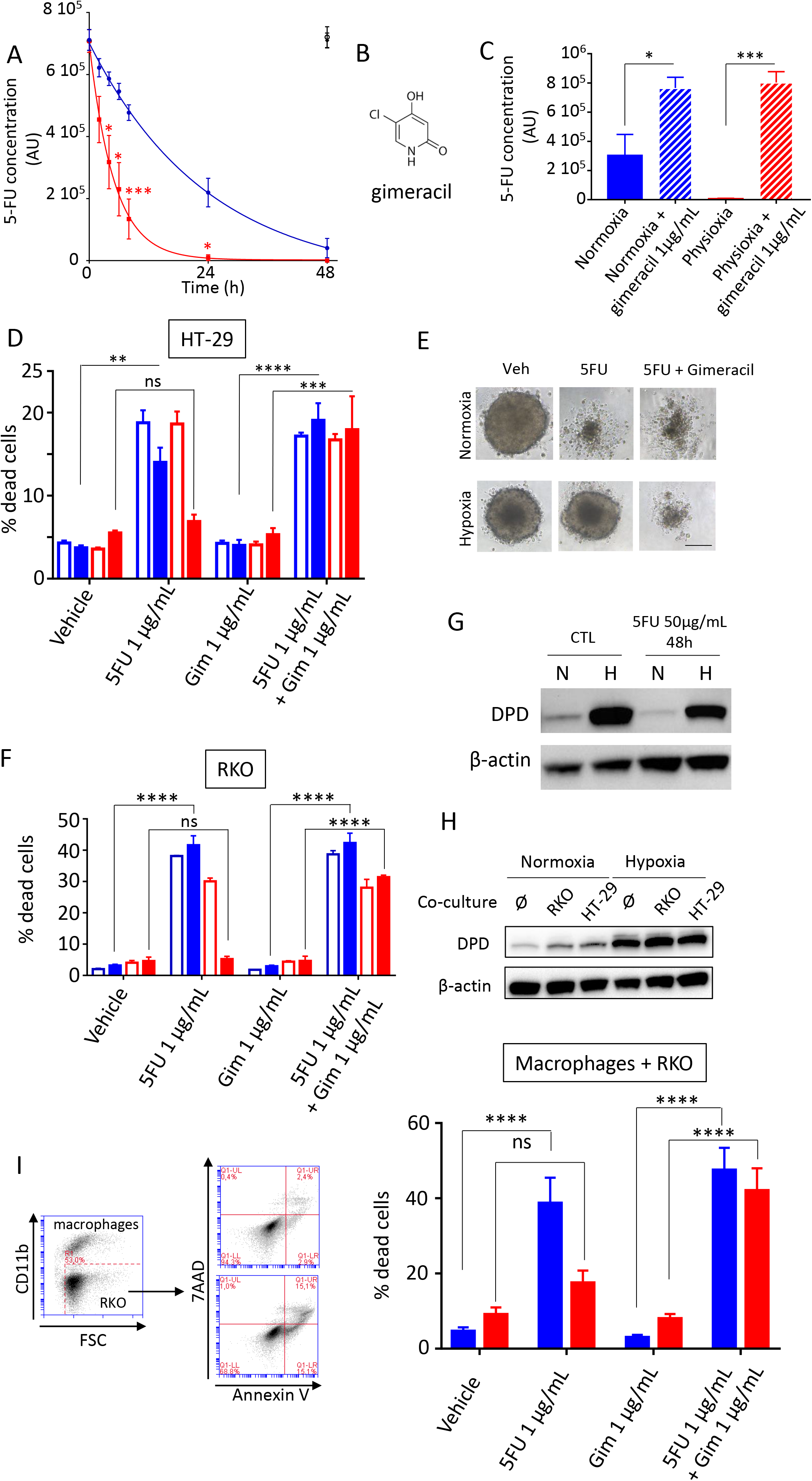
Chemoresistance to 5-FU is driven by DPD activity in macrophages. (A) Kinetics of 5-FU degradation by macrophages obtained by HPLC. 5-FU initial concentration was 1μg/mL. Normoxia appears in blue and hypoxia in red. 5-FU without macrophages was stable during the 48h period of study in normoxia (full black circle) and in hypoxia (empty black circle) (n=3). (B) Chemical structure of gimeracil a specific inhibitor of DPD. (C) 5-FU degradation due to DPD activity in macrophages inhibited by gimeracil. 5-FU initial concentration was 1 μg/mL. Normoxia appears in blue and hypoxia in red (n=3). (D) Induction of death in HT-29 in normoxia (blue) and in hypoxia (red) by non-conditioned medium (empty histogram) and conditioned medium (full histogram). Dead cells were defined as AnnexinV+ cells in flow cytometry. 5-FU was used at 1 μg/mL and gimeracil at 1 μg/mL (n=4). (E) Inhibition of growth and death induction in 3D tumoroïd of HT-29 cells in normoxia and hypoxia. Tumoroïds were exposed to CM(vehicle), CM(5-FU 1 μg/mL) and CM(5-FU 1 μg/mL + gimeracil 1 μg/mL). Picture were obtained with a phase contrast microscope, scale bar is 200 μm (n=8). (F) Induction of death in RKO in normoxia (blue) and in hypoxia (red) by non-conditioned medium (empty histogram) and conditioned medium (full histogram). Dead cells were defined as AnnexinV+ cells in flow cytometry. 5-FU was used at 1 μg/mL and gimeracil at 1 μg/mL (n=4). (G) Immunoblot of DPD expression in macrophages exposed to 5-FU at 50 μg/mL during 48 h (n=3). (H) Immunoblot of DPD expression in macrophages transwell co-cultured with HT-29 or RKO in normoxia and hypoxia (n=3). (I) Induction of death in RKO cells directly co-cultured with macrophages in normoxia (blue) and hypoxia (red). 5-FU was used at 1 μg/mL and gimeracil at 1 μg/mL. Dead cells were defined as CD11b-AnnexinV+ cells in flow cytometry. Gating strategy is represented on left panel. Dead cell quantification is represented on the right panel (n=4). Error bars represent mean ± sem, *p<0.05, **p<0.01, *** p<0.001, ****p<0.0001, ns=non significant

### Oxygen controls DPD expression in macrophages translationally through the cap-dependant protein synthesis molecular complex eIF4F^Hypoxic^

We discovered that a decreased oxygen concentration was able to increase the expression of DPD in human macrophages. In order to gain further insight into this, we carried out the transition of macrophages to various oxygen environments, to study the way DPD is controlled. We observed that DPD expression was inversely correlated to oxygen levels during the transitions (Figures 3A and 3B). The evolution of DPD expression was then analyzed with the help of a first-order differential equation (Figure 3C). This analysis revealed that the DPD half-life increased during the HN (Hypoxia to Normoxia) transition, demonstrating no increased degradation of DPD (Figure 3C). These results sugggested a synthesis control of DPD expression rather than a degradation control. We then demonstrated that hypoxic transitions were associated with the production of a functional DPD resulting in an increased dihydrouracil/uracil ratio in the extracellular medium (Figure 3D). Besides, we noted that profound hypoxic conditions (7 mmHg ~ 1% O2) provided the same induction of DPD overexpression as moderate hypoxic conditions (Figure 3E). As Hypoxic-induced factor HIF-1α is known to be stabilized during hypoxic transitions, so we checked whether its stabilization could be implicated in DPD over-expression. We found no induction of DPD synthesis due to HIF-1α stabilization in human macrophages (Figure 3F). We next sought to determine whether the expression of DPD is transcriptionally controlled when macrophages are exposed to low oxygen environments. To do so, we analyzed mRNA levels of oxygen-sensitive genes in macrophages (*VEGF-A*, *NDRG1*, *P4HA1*, *SLC2A1*) in the transition from normoxia to hypoxia or from hypoxia to normoxia. We discovered that the *DPYD* mRNA level did not present any significant variation of its level of expression compared to oxygen responsive genes (Figure 3G). We confirmed the absence of transcriptional control by inhibiting the synthesis of new mRNAs with actinomycin D and found no effect on DPD protein synthesis during a hypoxic transition (Figure 3H). This absence of correlation between the mRNA level and protein expression suggested a translational controlled mechanism. Recently, it has been demonstrated that the initial steps of protein synthesis such that as binding of the eukaryotic translational initiation factor E (eIF4E), part of the eIF4F^Normoxic^ initiation complex, to mRNA are repressed in hypoxia. Another complex, eIF4F^Hypoxic^, involving eIF4E2, RBM4 (RNA binding protein 4), and HIF-2α interacts with mRNA and mediates a selective cap-dependent protein synthesis in low oxygen environments (Ho et al., 2016; Uniacke et al., 2012). Genes containing a RNA hypoxic response element (rHRE) in their 3’UTR could bind to the HIF2α-RBM4-eIF4E2 complex part of eIF4F^Hypoxic^ (Uniacke et al., 2012). Therefore, we verified if *DPYD* presents such rHRE sequence on its 3’UTR and found effectively a trinucleotide CGG RBM4 binding motif (Figure 3I). Using these results, we depleted the expression of eIF4E and eIF4E2 in macrophages using specific siRNAs and found that hypoxia induced DPD synthesis is eIF4E2 dependent (Figure 3J). We also confirmed the involvement of HIF-2α in DPD synthesis in hypoxia (Figure 3K). These results demonstrated that DPD expression is controlled by the translation complex eIF4F^Hypoxic^ when oxygen decreases without involving a transcriptional regulation. Interestingly, the participation of HIF-2α in the eIF4F^Hypoxic^ complex is independent of its transcription factor activity (Uniacke et al., 2012). We have demonstrated that HIF-2α stabilization during hypoxia is the driving sensing signal that leads to the production of DPD in macrophages.

**Figure 3.**
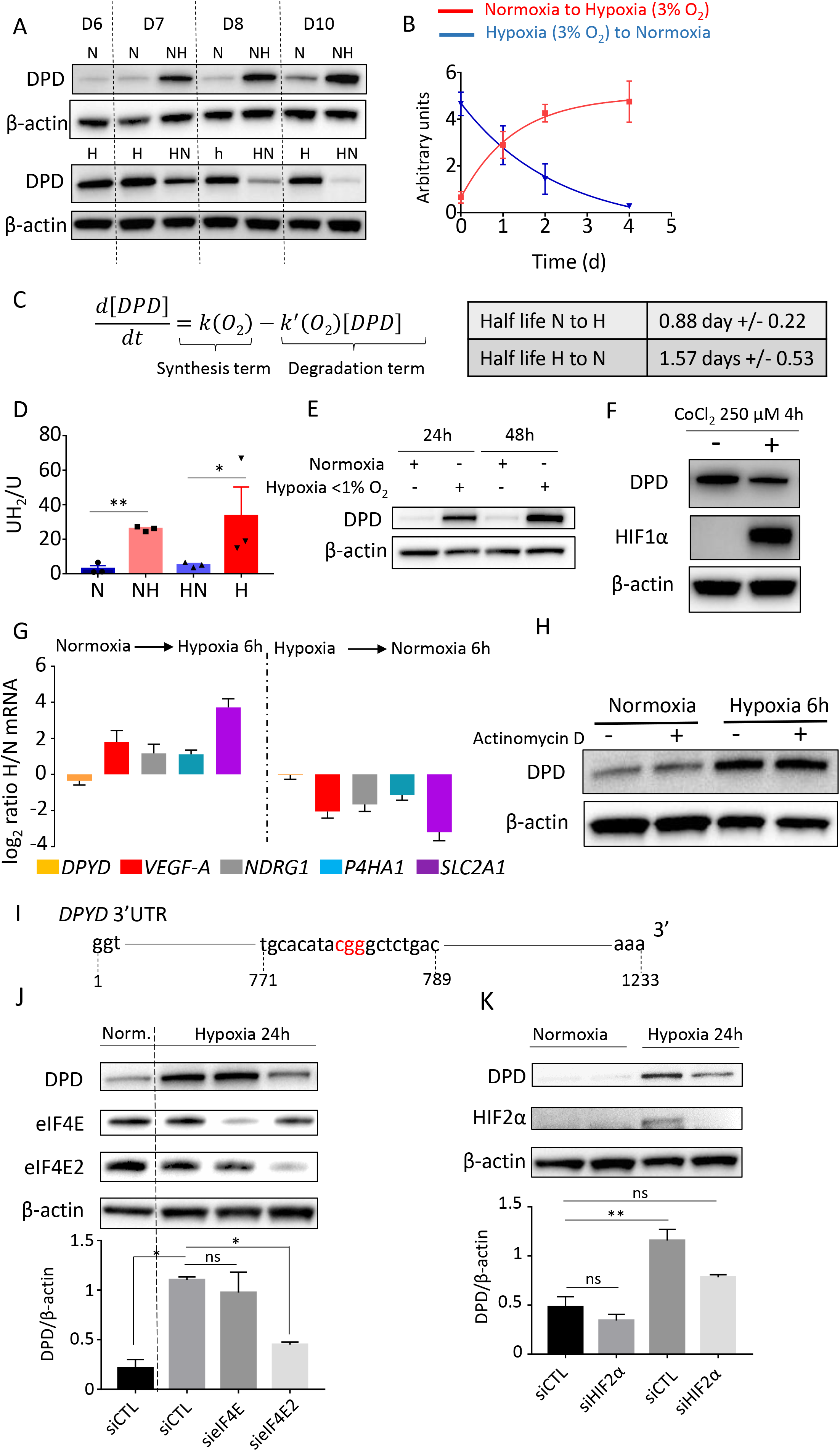
A translational mechanism controls DPD expression involving the eIF4F^Hypoxic^ complex. (A) Immunoblot of DPD expression in normoxia (N) and hypoxia (H) and during transition from normoxia to hypoxia (NH) and hypoxia to normoxia (HN) (n=4). (B) Quantification of the expression of DPD from immunoblots of figure 3A for NH and HN transitions (n=4). (C) Mathematical model fitting DPD expression curves from Figure 3B (left panel). DPD half-life extracted from the model (right panel). (D) DPD activity determined in the supernatant of macrophages cultured in normoxia and hypoxia for 24h. Macrophages were previously differentiated in normoxia and hypoxia permitting to study the activity of DPD following transitions (n=3). (E) Immunoblot of DPD expression in profound hypoxia for 24h and 48h (PO2 =7mmHg; n=4). (F) Immunoblot analysis of DPD and HIF-1α in macrophages exposed to CoCl2 250 μM for 4h. β-actin is used as control loading (n=3). (G) mRNA expression ratio NH/N and HN/H transitions determined by qPCR for the following genes *DPYD*, *VEGF-A*, *NDRG1*, *P4HA1* and *SLC2A1*. Macrophages were previously cultured in normoxia or hypoxia (n=3). (H) Immunoblot of DPD expression during NH (hypoxia PO_2_ =7mmHg) transition with macrophages exposed previously to Actinomycin D 1 μg/mL 20 minutes (n=3). (I) 3’UTR sequence of the DPYD gene containing a CGG RBM4 binding motif highlighted in red (Genecode transcript variant 1 ENST00000370192.7 https://genome.ucsc.edu/). (J) Immunoblot of DPD expression during NH transition under siRNA silencing of eIF4E2 and eIF4E (n=3). (K) Immunoblot of DPD expression during NH transition under siRNA silencing of HIF2α (n=3).

### TAMs in human colon cancer tissues harbor the principal component of DPD expression in tumors

We have demonstrated that DPD expression in macrophages confers a chemoresistance to 5-FU. In der to assess the clinical relevance of this result, we further determined the relative expression of DPD in various cellular populations found in colorectal tumors and colorectal liver metastasis. Based on immunohistochemical analysis of CRC tissues, it had been previously reported that cancer cells do not express DPD whereas normal cells, morphologically similar to macrophages, present a strong expression (Kamoshida et al., 2003). In order to appreciate the quantitative effect of this expression in macrophages in CRC, we analyzed the corresponding level of DPD expression in colorectal cancerous cells. We first used RNAseq analysis in various cancer cell lines from the Cancer Cell Line Encyclopedia (CCLE) and confirmed that the 59 cancer cell lines originating from colon cancer presented the lowest level of expression for *DPYD* suggesting a low expression pattern in these tumors (Figure 4A). This result confirmed what we had observed for three colon cancer cell lines RKO, HT-29 (Figure S1A), and Lovo (data not shown) and emphasized a preeminent role of macrophages in DPD induced chemoresistance in tumors. We further analyzed tissue samples from patients suffering from colorectal cancer with liver metastasis and found that the strongest expression of DPD was found in areas with CD68+ macrophages (Figure 4B). Tumor cells did not present a significant level of DPD expression in metastasis when compared to neighbouring TAMs (Figure 4B). Furthermore, in primary tumors, macrophages were also found to express the highest level of DPD, but no detectable expression was found in cancerous cells (Figure 4C). We further confirmed the exclusive expression of DPD by macrophages, by showing that strongly DPD+ cells were also CD68+ using an immunofluorescence co-expression analysis both in liver metastasis where (Figure 4E) and primary tumors (Figure 4F). Because CD68 has been found to be less specific than previously thought as a macrophage marker (Ruffell et al., 2012), we confirmed our results using the CD163 macrophage marker. We confirmed that DPD+ cells are CD163+ TAMs in liver metastasis and primary tumors (Figures S2A and S2B). Since macrophages can present various states of activation depending on their surrounding environment (Sica and Mantovani, 2012), we tested if DPD could be influenced by cytokines or growth factors. In this respect, we found that DPD expression in macrophages is not modulated according to the polarization state induced by IL-4, IL-6, IL-10, IL-13, IFNγ or GM-CSF, in contrast with the expression induced by hypoxia during the same period of time (Figure S2C). All these results indicate that DPD expression in colorectal cancers at the primary site and liver metastasis belong to macrophages, under the control of oxygen.

**Figure 4.**
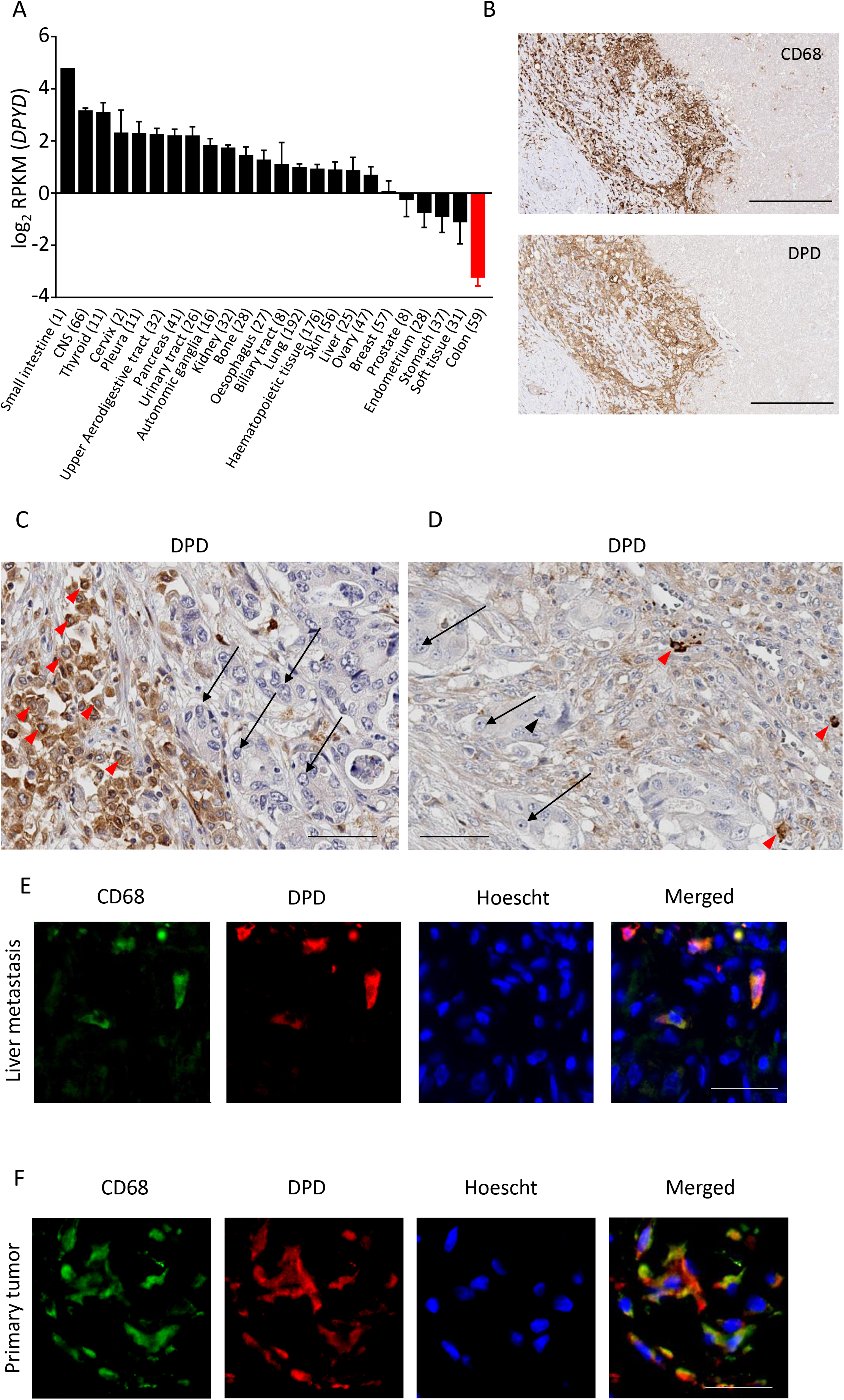
Macrophages harbor the main DPD expression in colorectal cancer. (A) RNAseq analysis of *DPYD* expression in various cancer cell lines from the Cancer Cell Line Encyclopedia (CCLE). Colon cancer cell lines are in red. (B) Immunochemistry analysis of CD68 (upper panel) and DPD (lower panel) expression in liver metastasis of colorectal cancer (n=15; scale bar = 200 μm). (C) Immunochemistry analysis of DPD expression in various cell populations in liver metastasis. Red arrowheads highlight macrophages, black arrows point to metastatic cancerous cells (n=15; scale bar = 60 μm). (D) Immunochemistry analysis of DPD expression in primary tumors. Red arrowheads are macrophages, black arrows point to cancer cells and the black arrowhead identifies a tripolar mitosis of a cancer cell (n=15; scale bar = 60 μm). (E) Immunofluorescence staining in liver metastatic tissues. CD68 is in green, DPD in red, nuclei are stained by Hoescht in blue (n=4; scale bar= 50 μm). (F) Immunofluorescence staining in primary tumors. CD68 is in green, DPD in red, nuclei are stained by Hoescht in blue (n=4; scale bar= 50 μm).

### Rodents’ macrophages do not express significant levels of DPD

In order to assess the generality of the oxygen control of DPD expression in macrophages we checked if this mechanism still holds in rodents. Surprisingly, we found that mice bone marrow derived macrophages (BMDM) do not express a significant level of DPD in normoxia or hypoxia despite the presence of the protein in mice liver (Figure 5A). No detectable level of *DPYD* mRNA was found in BMDM from BALB/c mice (data not shown). We discovered a similar result in the RAW264.7 mice macrophage cell line, where no protein (Figure 5B) or mRNA of *DPYD* was found (data not shown). Similarly, we also found that chemically mobilized peritoneal macrophages from adult female Fischer rats do not express DPD (data not shown). We then used open data sets from microarrays to compare the expression of *DPYD* mRNA levels in humans to those in mice and found that *DPYD* presents the highest levels of expression in human monocyte/macrophages from various anatomical compartments, contrary to what was found in mice where macrophages expressed few mRNA *DPYD* molecules compared to other cellular compartments (Figure 5C). This result suggests repression of mRNA synthesis in mice macrophages. We confirmed the epigenetic control of gene expression using 5-aza-2’deoxycytidine (decitabine) a DNA hypo-methylation agent. Indeed, decitabine led to a strong increase in m*DPYD* mRNA level of expression in treated RAW macrophages compared to non-treated cells (Figure 5D). This result suggests that the methylation of the *DPYD* promoter gene in mice is, at least in part, responsible for its repressed expression.

**Figure 5.**
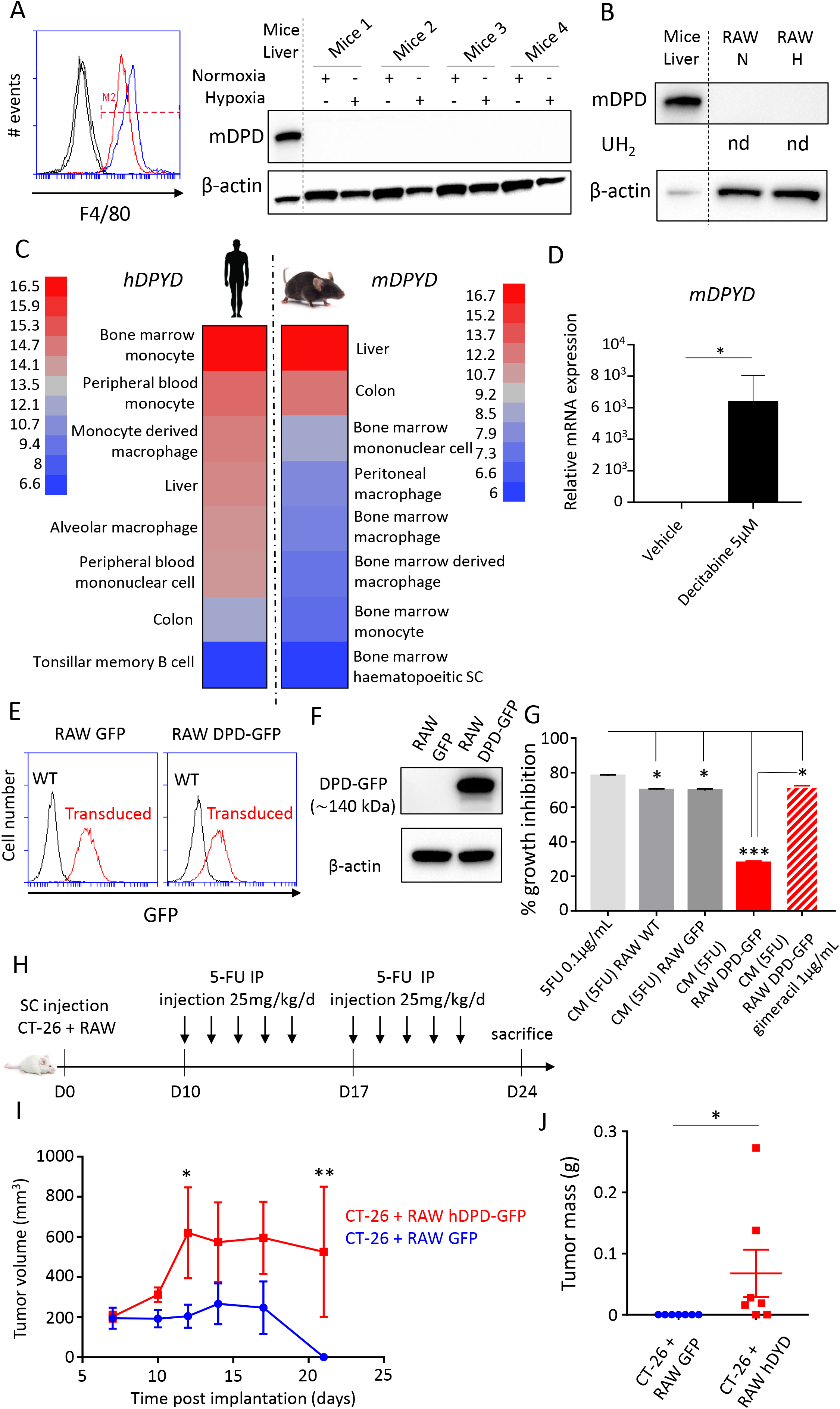
Rodents’ macrophages do not express DPD and transduced human DPD in mice macrophages leads to 5-FU chemoresistance *in vivo*. (A) BMDM were differentiated in normoxia and hypoxia. F4/80 was studied by flow cytometry (left panel) and mDPD expression by immunoblot (n=5). Mice liver was used as a positive control for DPD. (B) RAW264.7 macrophages were cultivated in normoxia and in hypoxia. mDPD expression was studied by immunoblot. No production of dihydrouracil was found in RAW supernatant using HPLC (nd=non detected). (C) Microarray analysis of *DPYD* mRNA expression in monocytes/macrophage populations in mice and humans. In each group the highest and lowest level of expression was used to scale the heat map. (D) m*DPYD* mRNA level of expression in RAW macrophages exposed to decitabine at 5μM during 24h (n=3). (E) Transduced GFP and hDPD-GFP proteins levels of expression analyzed by flow cytometry. (F) Immunoblot of RAW macrophages transduced to express GFP and hDPD-GFP (G) Growth inhibition of CT-26, after 48h, under the presence of CM containing 5-FU 0.1μg/mL exposed to macrophages WT, expressing GFP or hDPD-GFP for 24h. Gimeracil was used to block DPD activity at 1 μg/mL. (H) Tumor assay was performed on female Balb/c mice of 7 weeks. 106 CT-26 and 106 RAW were implanted subcutaneously. After ten days daily bolus of 5-FU 25 mg/kg were injected intraperitoneally according to the timeline represented. (I) Tumors growth was followed during the protocol (n=7 in each group). (J) Tumors weight was determined at day 24 (n=7 in each group).

### Transduced human *DPYD* in mice macrophages leads to 5-FU chemoresistance *in vivo*

In order to obtain a mice model mimicking the human DPD expression in macrophages, we transduced the human *DPYD* gene incorporated into a lentivirus to obtain “DPD-humanized” mice macrophages (Figures 5E and 5F). CM of mice macrophages expressing DPD were able to confer chemoresistance to CT-26 (a mice colon cancer cell line that does not express DPD) demonstrating the functionality of the transduced *DPYD* (Figure 5G). We also discovered that wild-type macrophages are associated with weak protection toward 5-FU compared to macrophages expressing DPD, demonstrating that even if other chemoresistance mechanisms are associated with TAMs DPD-induced chemoresistance is probably the more efficient one (Figure 5G). In order to further confirm the relationship between DPD expression in macrophages and chemoresistance in colorectal cancers, a tumor assay in mice was performed. CT-26 and RAW macrophages expressing or not human DPD were implanted into flanks of BALB/c mice. Ten days after the implantation, 5-FU at 25mg/kg was injected intraperitoneally during five days for two consecutive weeks (Figure 5H). We confirmed that tumors harboring macrophages expressing DPD were more resistant to 5-FU than the control tumors with wild-type macrophages (Figures 5I and 5J), indicating that DPD expression in TAMs promotes chemoresistance *in vivo*.

## Discussion

In recent years, the function of the immune system has become a key element in understanding tumor interaction with the surrounding healthy tissues as well as a provider of new therapeutic strategies. The tumor immune microenvironment (TIME) is composed of various types of immune cells; nevertheless, TAMs usually represent quantitatively the largest population found in solid cancers. TAMs are involved in tumor growth, immune evasion, neoangiogenesis and treatment resistance. Using depletion methods, a large number of studies have reported an increased chemo-sensitivity when macrophages are removed from the tumor (Ruffell and Coussens, 2015). Futhermore, co-culture studies have revealed a macrophage-mediated resistance to various anti-cancer drugs such as paclitaxel, doxorubicin, etoposide or gemcitabine (Mitchem et al., 2013; Shree et al., 2011). Specifically, depletion of MHCII^lo^ TAMs leads to an increased sensitivity to taxol-induced DNA damage and apoptosis (Olson et al., 2017). Another key point is that TAMs were mainly found in hypoxic areas where they could further favor hypoxia by secreting VEGF-A, leading to the formation of an abnormal vasculature (Murdoch et al., 2008). Accordingly, mechanisms involving macrophages-induced chemoresistance relied on an indirect effect due to the secretions by macrophages such as pyrimidine nucleosides (deoxycytidine) inhibiting gemcitabine induction of apoptosis in pancreatic ductal adenocarcinoma (Halbrook et al., 2019). More specifically in CRC, the implication of macrophages in chemoresistance has also been suggested based on *in vitro* and *in vivo* studies. The mechanisms proposed were diverse but involved indirect secreted factors. For example, it has been proposed that IL-6 secreted by macrophages could stimulate STAT3 in cancerous cells inducing the inhibition of the RAB22A/BCL2 pathway through miR-204-5p expression, thereby leading to chemoresistance to 5-FU (Yin et al., 2017). Similarly the secretion of putrescin by macrophages, a member of the polyamine family, was shown to suppress the JNK/Caspase 3 pathway in cancerous cells, providing a protection against 5-FU (Zhang et al., 2016).

To order to understand the involvement of TAM in chemoresistance in CRC, we designed the present study to incorporate oxygen concentration as a key environmental parameter. We had previously reported that the oxygen disponibility in macrophages’ environment greatly influences their immune functions such as their ability to clear apoptotic cells (Court et al., 2017). As colon tissues are naturally exposed to levels of oxygen that are usually lower than 5% O_2_ (Keeley and Mann, 2018) with values that could reach even lower (<1% O_2_) in tumors, oxygen appeared as a fundamental parameter to understand macrophage involvement in chemoresistance. We first verified if a secreted factor by human macrophages could provide a chemoresistance and finally found a direct effect of hypoxic macrophages on 5-FU (Figures 1A and 1B). A molecular analysis revealed that the dihydropyrimidine deshydrogenase (DPD), an enzyme of the pyrimidine catabolism pathway, was over-expressed in hypoxic human macrophages providing a direct chemoresistance mechanism (Figures 1C and 2). We found that DPD was part of the protein synthesis response to hypoxia in macrophages under the control of the translation initiating complex eIF4F^Hypoxic^ (Figure 6), notably composed of HIF-2α that is stabilized in hypoxia but acts in this complex independently of its transcription factor activity (Uniacke et al., 2012). Accordingly, we found that this mechanism was independent of polarization states, as various polarizing factors were unable to modify DPD expression contrary to hypoxia (Figure S2B). This result demonstrated the robustness of the oxygen control of DPD expression, irrespective of the polarization states. As TAM populations have been demonstrated to be highly heterogeneous according to their phenotype, which does not fit the classical M1/M2 classification (Aras and Zaidi, 2017), the universality of the mechanism identified in human macrophages assured its relevance in colorectal cancers (Figures 4B to 4F and S2A).

**Figure 6.**
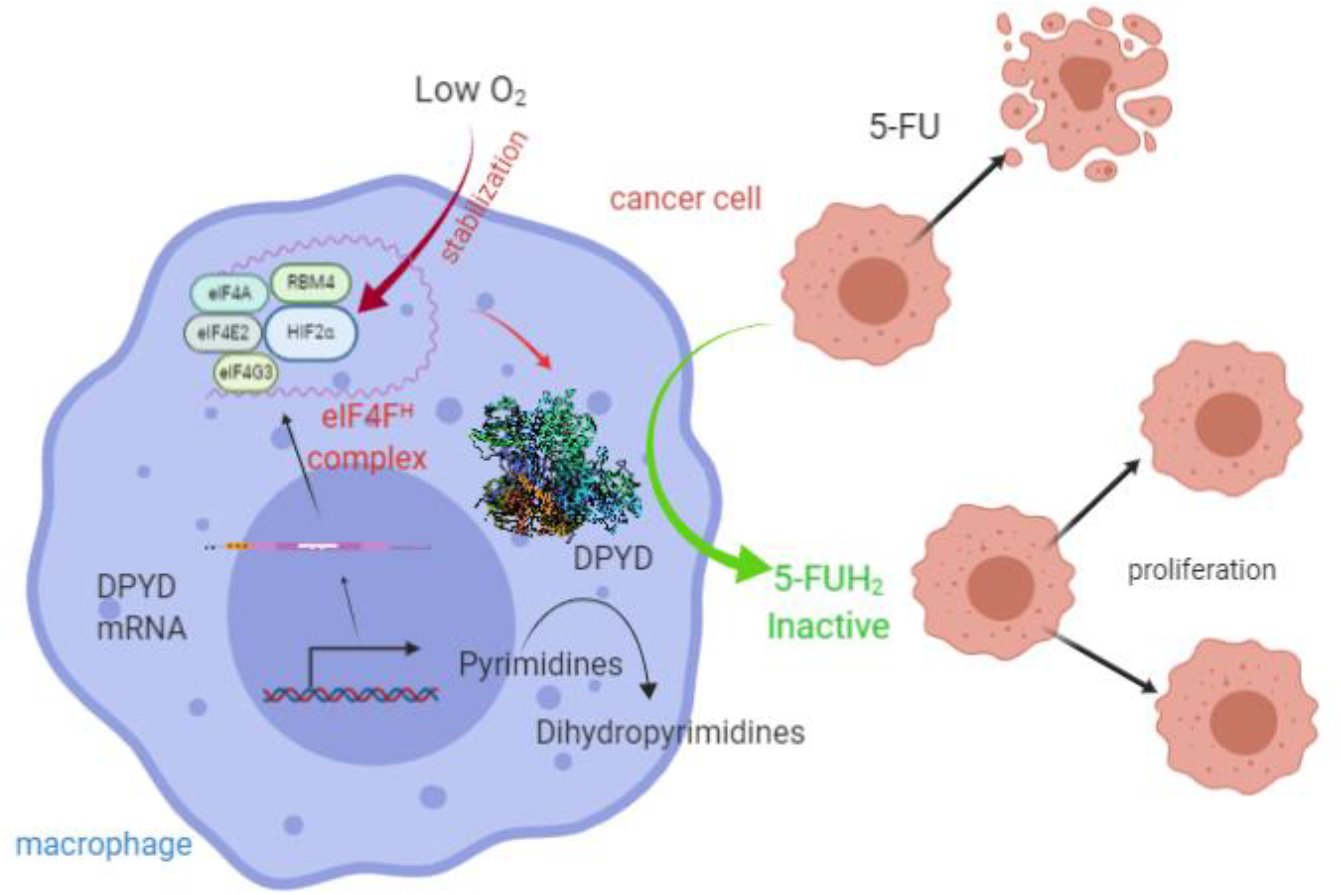
DPD expression in macrophages confers a chemoresistance to 5-FU under the control of oxygen. Schematic of the chemoresistance mechanism due to DPD expression in hypoxic macrophages under the control of the eIF4F^Hypoxic^ complex. 5-FU degradation is mainly performed by macrophages in the tumor microenvironement providing a protection for cancer cells, free to proliferate.

DPD expression in macrophages seemed to be particularly relevant in CRC where cancerous cells present a low expression level of the protein in primary tumors as well as in liver metastasis (Figures 4A, 4C and 4D), assuring macrophages to be mainly responsible for the tumor tissue degradation of 5-FU. This general feature seems to rely on the epigenetic control of DPD expression in cancer cells. Indeed, many colon cancer cells lines have been noted to harbor a histone H3K27me3 mark that blocks the fixation of the transcription factor PU.1, leading to the inhibition of DPD mRNA transcription (Wu et al., 2016).

We further found that macrophages in rodents do not express DPD as it is transcriptionally repressed. This finding forced us to re-evaluate previous *in vivo* models that are used to assess the involvement of macrophages in CRC. Indeed, we have shown the importance of DPD activity in macrophages and found that it represents the main quantitative source of degradation of 5-FU in human colorectal tumors. In order to demonstrate the relevance of this mechanism to chemoresistance, we designed an *in vivo* model using mice macrophages expressing DPD. We validated the importance of DPD expression in TAMs leading to chemoresistance to 5-FU. Supporting these results, previous clinical studies have suggested that DPD mRNA expression is a marker of chemoresistance in CRC. These studies usually assumed that DPD expression is mainly caused by cancerous cells. Nevertheless, the mRNA level in the tumor is correlated with a low response to 5-FU confirming its relevance (Ichikawa et al., 2003; Salonga et al., 2000; Shirota et al., 2002; Soong et al., 2008). The results we have obtained in our study suggest that the main prognostic factor for 5-FU response is DPD expression in macrophages located in tumors and liver metastasis. That expression is notably important in the invasive front, where TAMs seem to concentrate (Pinto et al., 2019). Furthermore, the invasion front is known to be a hypoxic area in CRCs (Righi et al., 2015). Since the mechanism identified in this study relied on quantitative expression of DPD by macrophages, the assessment of the spatial heterogeneity of DPD expression will be necessary to stratify patients in various response groups for chemotherapy (Marusyk et al., 2020). Thus this study constitutes an important advance in the understanding of the role of tumor immune environment in chemoresistance to 5-FU in CRC. Finally, further translational and clinical studies are needed to confirm the clinical relevance of these findings and develop new predictive markers of response to 5-FU-based treatments in order to improve colorectal patients’ care.

## MATERIALS AND METHODS

### Cell culture

RAW264.7, CT-26, RKO, HT-29, Lovo were purchased from ATCC. RAW were maintained in high-glucose DMEM (Gibco) supplemented with 10% FBS (Gibco) at 37°C, CT-26 and RKO were maintained in RPMI (Gibco) supplemented with 10% FBS (Gibco) at 37°C, HT-29 were maintained in McCoy’s medium (Gibco) supplemented with 10% FBS (Gibco) at 37°C and Lovo were maintained in F12-K supplemented with 10% FBS (Gibco) at 37°C. All cells were routinely tested for mycoplasma contamination using MycoAlert detection kit (Lonza). All cells have been used in the following year of their reception.

### Human samples

Human blood samples from healthy de-identified donors are obtained from EFS (French national blood service) as part of an authorized protocol (CODECOH DC-2018–3114). Donors gave signed consent for use of their blood in this study. Tumor samples were obtained from the department of pathology of the university hospital of Grenoble as part of a declared sample collection AC-2014-2949. Patient selection criteria were a diagnostic of colorectal adenocarcinoma and tissue samples availability for the primary tumor and liver metastasis. Clinical characteristics of the patients are reported in the table. All patients gave their signed consent for this study as part of an authorized protocol (INDS MR0709280220).

### Animals

8 weeks old Balb/c female mice were obtained from Charles River. Animals were housed and bred at « Plateforme de Haute Technologie Animale (PHTA) » UGA core facility (Grenoble, France), EU0197, Agreement C38-51610006, under specific pathogen–free conditions, temperature-controlled environment with a 12-h light/dark cycle and ad libitum access to water and diet. Animal housing and procedures were conducted in accordance with the recommendations from the Direction des Services Vétérinaires, Ministry of Agriculture of France, according to European Communities Council Directive 2010/63/EU and according to recommendations for health monitoring from the Federation of European Laboratory Animal Science Associations. Protocols involving animals were reviewed by the local ethic committee “Comité d’Ethique pour l’Expérimentation Animale no.#12, Cometh-Grenoble” and approved by the Ministry of Research under the authorization number (January 2020) APAFIS#22660-2019103110209599.

### Human macrophage differentiation from monocytes

Monocytes were isolated from leukoreduction system chambers of healthy EFS donors using differential centrifugation (Histopaque 1077, Sigma) to obtain PBMCs. CD14^+^ microbeads (Miltenyi Biotec) were used to select monocytes according to the manufacturer’s instructions. Monocytes were plated in RPMI (Life Technologies) supplemented with 10% SAB (Sigma), 10 mM HEPES (Life Technologies), MEM Non-essential amino acids (Life Technologies) and 25 ng/ml M-CSF (Miltenyi Biotec). Differentiation was obtained after 6 days of culture. Hypoxic cultures were performed in the HypoxyLab chamber authorizing an oxygen partial pressure control (Oxford Optronix, UK).

### Bone-Marrow Derived Macrophage Differentiation

Bone marrow was extracted from the femurs of Balb/c mice in RPMI and then filtered by a 70μm cell strainer. Cells were washed in RPMI and then cultured in RPMI (Gibco) supplemented with 10% of FBS (Gibco) and mice M-CSF at 25 ng/mL (Miltenyi Biotec) for 6 days. Medium was refreshed at day 3. Differentiation was assessed by flow cytometry analysis of surface F4/80 expression.

### Conditioned medium

Macrophages at 1 × 10^6^ cells per well in 12-wells plates were cultured with RPMI supplemented with 10% SAB with DMSO (vehicle), Gimeracil 1 μg/ml, 5-FU 0.1 μg/ml or 1 μg/ml or 5-FU 0.1 μg/ml or 1 μg/ml with Gimeracil 1 μg/ml during 24h in normoxia and hypoxia. The CM was added to HT-29 and RKO, at 3 × 10^5^ cells per well in 12-wells plates for 48 hours. Then cells were collected, counted and the mortality rate assessed by flow cytometry (Annexin V and 7-AAD).

### RNAi

Small interfering RNA (siRNA) (GE Dharmacon) was transfected at a final concentration of 50 nM using Lipofectamine (RNAiMAx, Life Technologies).

### Expression of human DPD in mice macrophages

RAW264.7 were transduced using lentivirus particles with human DPD (mGFP-tagged) inserted in a pLenti-C-mGFP-P2A-Puro plasmid (Origen Technologies, Rockville, US). Control RAW were obtained using the lentivirus particles containing the plasmid without the DPD ORF sequence (pLenti-C-mGFP-P2A-Puro).

### RNA isolation and qPCR analysis for gene expression

Cells were directly lysed and RNA was extracted using the NucleoSpin RNA kit components (Macherey Nagel) according to the manufacturer’s instructions. Reverse transcription was performed using the iScript Ready-to-use cDNA supermix components (Biorad). qPCR was then performed with the iTaq universal SYBR green supermix components (Biorad) on a CFX96 (Biorad). Quantification was performed using the ΔΔCt method.

### Immunoblotting

Cells were lysed in RIPA buffer supplemented with antiprotease inhibitors (AEBSF 4 mM, Pepstatine A 1 mM and Leupeptine 0.4 mM; Sigma Aldrich) and HIF-hydroxylase inhibitor (DMOG 1 mM, Sigma Aldrich). Proteins were quantified by BCA assay (ThermoFischer) and 15 μg of total protein was run on SDS-PAGE gels. Proteins were transferred from SDS-PAGE gels to PVDF membrane (Biorad), blocked with TBS supplemented with 5% milk then incubated with primary antibody at 1μg/mL overnight 4°C. After washing with TBS, the membrane was incubated with a horseradish peroxidase-conjugated secondary antibody (Jackson Immunoresearch). Signal was detected by chemoluminiscence (Chemi-Doc Imaging System, Bio-Rad) after exposition to West Pico ECL (ThermoFischer).

### Proteomics

Cells are directly lysed in Laemmli buffer and prepared and analyzed as previously described (Court et al., 2017). Briefly, the protein equivalent of 300 000 cells for each sample was loaded on NuPAGE Bis-Tris 4–12% acrylamide gels (Life Technologies). Electrophoretic migration was controlled to allow each protein sample to be split into six gel bands. Gels were stained with R-250 Coomassie blue (Bio-Rad) before excising protein bands. Gel slices were washed then dehydrated with 100% acetonitrile (Merck Millipore), incubated with 10mM DTT (Dithiothreitol, Merck Millipore), followed by 55mM iodoacetamide (Merck Millipore) in the dark. Alkylation was stopped by adding 10mM DTT in 25mM ammonium bicarbonate. Proteins were digested overnight at 37 °C with Trypsin/Lys-C Mix (Promega, Charbonnières, France) according to manufacturer’s instructions. After extraction, fractions were pooled, dried and stored at −80 °C until further analysis. The dried extracted peptides were resuspendedand analyzed by online nano-LC (Ultimate 3000, Thermo Scientific) directly linked to an impact IITM Hybrid Quadrupole Time of-Flight (QTOF) instrument fitted with a CaptiveSpray ion source (Bruker Daltonics, Bremen, Germany). All data were analyzed using Max-Quant software (version 1.5.2.8) and the Andromeda search engine. The false discovery rate (FDR) was set to 1% for both proteins and peptides, and a minimum length of seven amino acids was set. MaxQuant scores peptide identifications based on a search with an initial permissible mass deviation for the precursor ion of up to 0.07 Da after time-dependent recalibration of the precursor masses. Fragment mass deviation was allowed up to 40 ppm. The Andromeda search engine was used to match MS/MS spectra against the Uniprot human database (https://www.uniprot.org/). Enzyme specificity was set as C-terminal to Arg and Lys, cleavage at proline bonds and a maximum of two missed cleavages are allowed. Carbamidomethylation of cysteine was selected as a fixed modification, whereas N-terminal protein acetylation and methionine oxidation were selected as variable modifications. The “match between runs” feature of MaxQuant was used to transfer identification information to other LC-MS/MS runs based on ion masses and retention times (maximum deviation 0.7 min); this feature was also used in quantification experiments. Quantifications were performed using the label free algorithms. A minimum peptide ratio counts of two and at least one “razor peptide” were required for quantification. The LFQ metric was used to perform relative quantification between proteins identified in different biological conditions, protein intensities were normalized based on the MaxQuant “protein group.txt” output (reflecting a normalized protein quantity deduced from all peptide intensity values). Potential contaminants and reverse proteins were strictly excluded from further analysis. Three analytical replicates from three independent biological samples (donors) were analyzed for each normoxic and hypoxic conditions. Missing values were deduced from a normal distribution (width: 0.3; down shift: 1.8) using the Perseus (version 1.5.5.3) post data acquisition package contained in MaxQuant (www.maxquant.org). Data were further analyzed using JMP software (v.13.0.0, SAS Institute Inc.). Proteins were classed according to the paired Welch test difference (difference between the mean value for triplicate MS/MS analyses for the two conditions compared), and the median fold-change between the two conditions compared.

### Immunochemistry

3 μm thick consecutive tissue sections were prepared from formalin-fixed and paraffin-embedded tissues. Deparaffinization, rehydratation, antigen retrieval and peroxidase blocking were performed on a fully automated system BENCHMARK ULTRA (Roche) according to manufacturer recommendations. The sections were incubated with the following primary antibodies: anti-CD3 clone 2GV6 (Roche), anti-CD68 clone Kp1 (Dako) and anti-DPD (ThermoFischer). Revelation was performed using the Ultraview DAB revelation kit (Roche). Nuclei were counterstained with hematoxylin solution (Dako). Images were captured using an APERIO ATS scanner (Leica).

### Immunofluorescence

Formalin fixed and paraffin-embedded human tissue samples were sectioned at 3μm thickness. Samples were de-paraffinized and hydrated by xylene and decreasing concentrations of ethanol baths, respectively. Antigen retrieval was achieved using IHC-TekTM Epitope Retrieval Steamer Set (IW-1102, IHCworld, USA) in IHC-TekTM Epitope Retrieval (IW-1100, IHCworld, USA) for 40 minutes. Nonspecific binding sites were blocked by 1% BSA in PBS. Samples were incubated with the primary antibodies: Monoclonal Mouse Anti-Human CD68 clone PG-M1 at 0.4 μg/mL, Mouse Anti-Human CD163 clone EDHu-1 at 10 μg/mL and DPD polyclonal antibody at 3 μg/mL for 1 hour at room temperature followed by an incubation of secondary antibodies: Alexa Fluor 488 goat anti-mouse IgG (H+L) and Alexa Fluor 546 goat anti-rabbit IgG (H+L) both at 4 μg/mL for 30 minutes at room temperature. Nuclei were stained by Hoechst 33342 at 5 μg/mL for 5 minutes at room temperature. Images were captured under 20X magnification using ApoTome microscope (Carl Zeiss, Germany) equipped with a camera AxioCam MRm and collected by AxioVision software and analyzed using ImageJ software.

### Flow cytometry

Flow cytometry data was acquired on an Accuri C6 (BD) flow cytometer. The reagents used were: AnnexinV-FITC, mouse anti-F4/80-PE clone REA126 and human anti-CD14-FITC from Miltenyi Biotech and 7-AAD staining solution from BD Pharmingen. Doublet cells were gated out by comparing forward scatter signal height (FSC-H) and area (FSC-A). Dead cells were excluded based on 7AAD or FSC/SSC profile. A least 10,000 events were collected in the analysis gate. Median fluorescence intensity (MFI) was determined using Accuri C6 software (BD).

### Cancer cell line mRNA *DPYD* expression

Cancer cell line gene expression data were collected from the Cancer Cell Line Encyclopedia (CCLE) (https://portals.broadinstitute.org/ccle). Briefly, sequencing was performed on the Illumina HiSeq 2000 or HiSeq 2500 instruments, with sequence coverage of no less than 100 million paired 101 nucleotides-long reads per sample. RNaseq reads were aligned to the B37 version of human genome using TopHat version 1.4. Gene and exon-level RPKM values were calculated using pipeline developed for the GTEx project (Barretina et al., 2012; Tsherniak et al., 2017). The cell lines were classified based on their tissue of origin resulting in 24 different groups. The number of cell lines in each group is indicated. The histogram plot was generated using the JMP software (SAS).

### Mice and human mRNA *DPYD* expression analysis

Genenvestigator 7.5.0 (https://genevestigator.com/gv/) is a search engine that summarizes data sets in metaprofiles. GENEVESTIGATOR^®^ integrates manually curated and quality-controlled gene expression data from public repositories (Hruz et al., 2008). In this study, the Condition Tools Search was used to obtain DPD mRNA levels obtained from human (*Homo sapiens*) and mice (*Mus musculus*) in various anatomical parts. Mean of logarithmic level of expression obtained from AFFIMETRIX expression microarrays was used to generate a cell plot from selected results related to macrophages and monocytes. The lowest and the highest expression results in both series (human and mice) were used to evaluate the expression level in the dataset. Cell plot was generated using the JMP software (SAS).

### DPD activity measurements

Since DPD is involved in the hydrogenation of uracil (U) into dihydrouracil (UH2), DPD activity was indirectly evaluated in the cell culture supernatants by determining U and its metabolite UH2 concentrations followed by the calculation of the UH2/U ratio. These analyses were performed in the Pharmacology Laboratory of Institut Claudius-Regaud (France) using an HPLC system composed of Alliance 2695 and diode array detector 2996 (Waters), according to a previously described method (Thomas et al., 2016). Uracil (U), Dihydrouracil (UH2), 5-FluoroUracil (5-FU), Ammonium sulfate 99%, Acetonitrile (ACN) gradient chromasolv for HPLC and 2-propanol were purchased from Sigma. Ethyl acetate Scharlau was of HPLC grade and purchased from ICS (Lapeyrouse-Fossat, France). The water used was of Milli-Q Advantage A10 as well as MultiScreen-HV 96-well Plates (Merck Millipore). Calibration ranges were 3.125 – 200 ng/ml for U and 25 - 500 ng/ml for UH2 and 5-FU (5μg/ml) was used as an internal standard. For the 5-FU kinetics experiments, no internal standard was added to the samples and the amount of 5-FU in the supernatant was quantified by the peak area corresponding to 5-FU.

### Quantification and statistical analysis

Statistics were performed using Graph Pad Prism 7 (Graph Pad Software Inc). When two groups were compared, we used a two-tailed student’s t test when variable presented a normal distribution or a Mann-Withney non parametric test otherwise. When more than two groups were compared, we used a one-way ANOVA analysis with Tukey post hoc test. All error bars represent mean with standard error of the mean, all group numbers and explanation of significant values are presented within the figure legends.

### Data availabilty

All MS proteomics data were deposited on the Proteome-Xchange Consortium website (http://proteomecentral.proteomexchange.org) via the PRIDE partner repository, data set identifier: PXD006354.

## Acknowledgments

AM is supported by the ATIP/Avenir Young group leader program (Inserm and La ligue nationale contre le cancer). KG is supported by the ITN (International Training Network) Phys2Biomed project which is funded from the European Union’s Horizon 2020 research and innovation programme under the Marie Skłodowska-Curie grant agreement No 812772.The authors would like to thank Xavier Fonrose, Edwige Col and Floriane Laurent for their technical assistance. The authors thank the zootechnicians of PHTA facility for animal housing and care.

## Authors’ contributions

AM conceived and designed the study. AM planned and guided the research and wrote the manuscript. MM, MC, KG, and AM performed experiments. MM, MC, KG, MHL, TD, GR, AM analyzed and interpreted data. SM,FT performed DPD activity measurements. MHL performed the histopathological analysis. AM supervised the work carried out in this study and raised funding.

## Supplementary materials

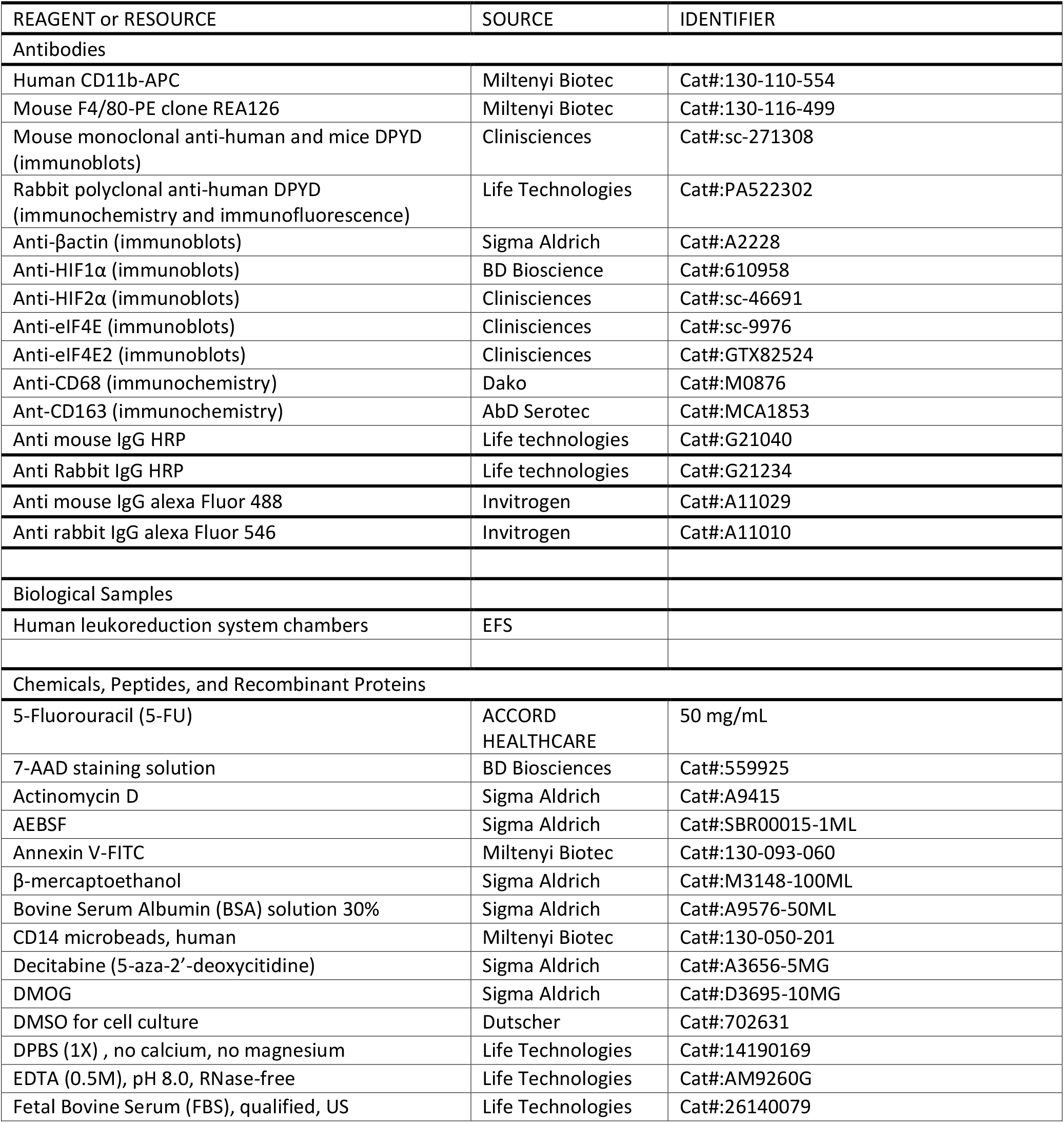
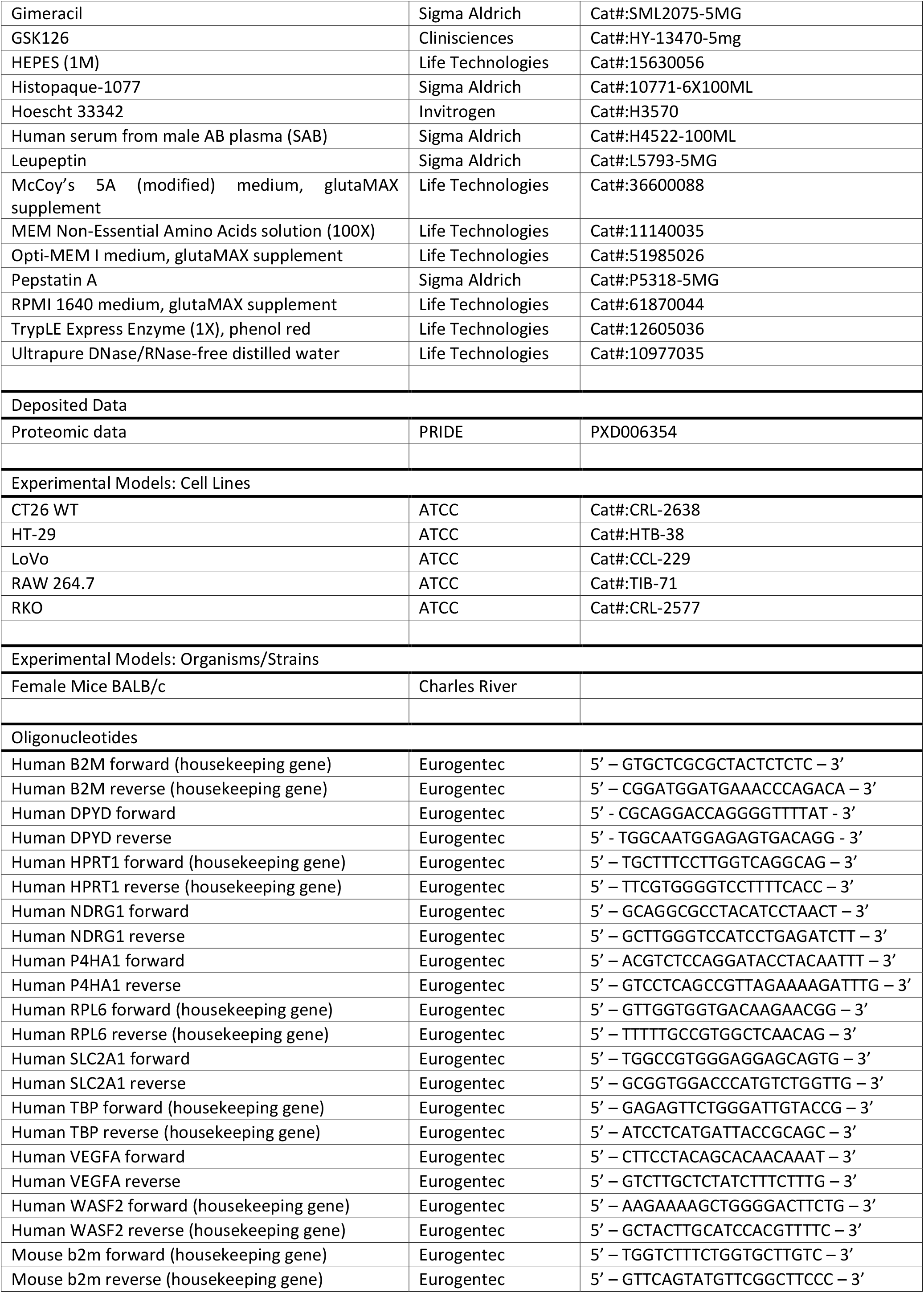
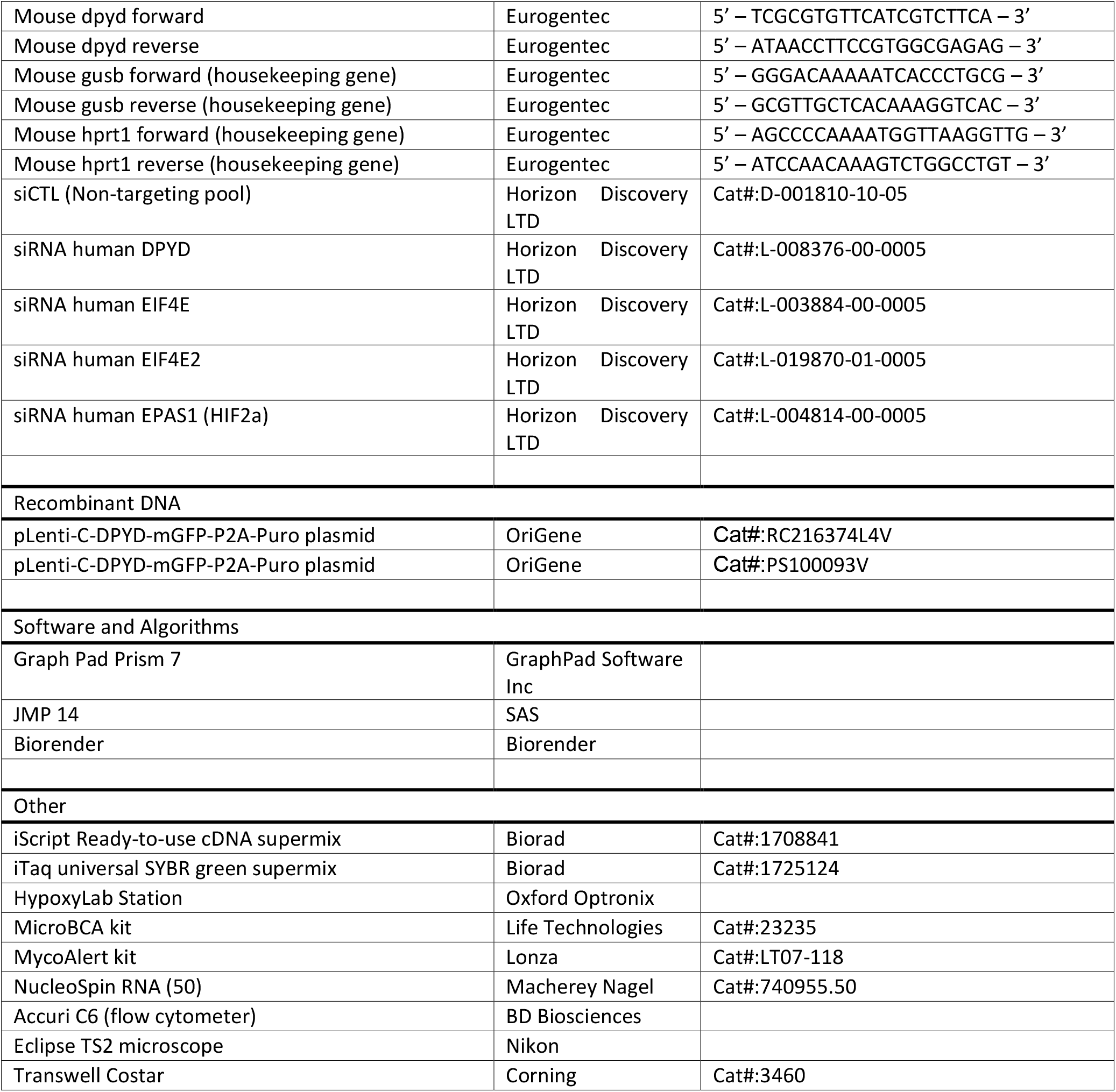

## Supplemental Figures

**Figure S1.**
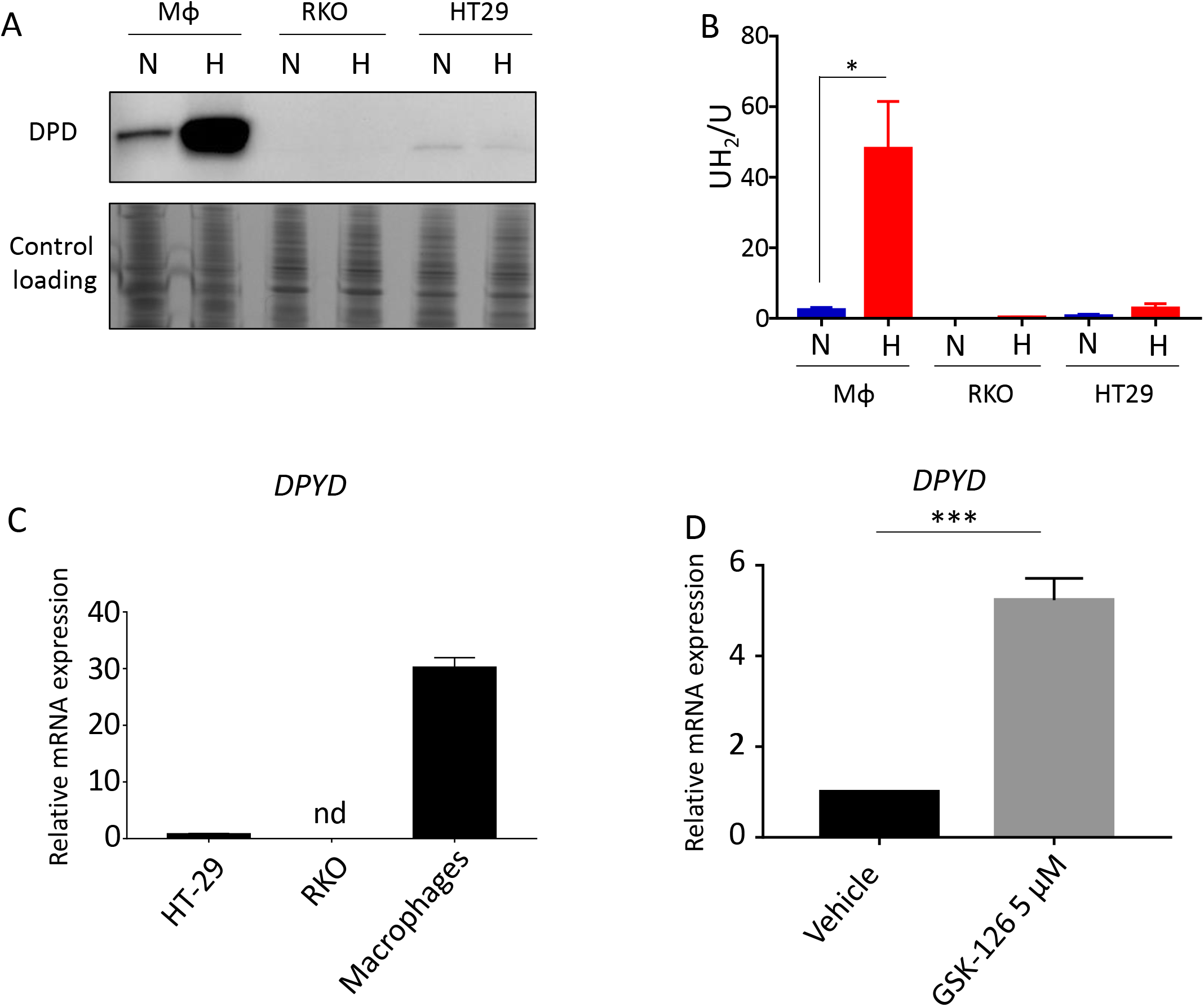
(A) Immunoblot analysis of DPD expression in human macrophages, RKO and HT-29 cells in normoxia and hypoxia (n=3). (B) Dihydrouracil to uracil ratio in the supernatant of human macrophages, RKO and HT-29 cells in normoxia and hypoxia measured by HPLC (n=3). (C) qPCR analysis of *DPYD* mRNA in human macrophages, RKO and HT-29 cells (n=3). nd=non detected (D) qPCR analysis of *DPYD* mRNA in RKO cells exposed to GSK-126 (an EZH2 inhibitor) at 5 μM during 96h (n=3)

**Figure S2.**
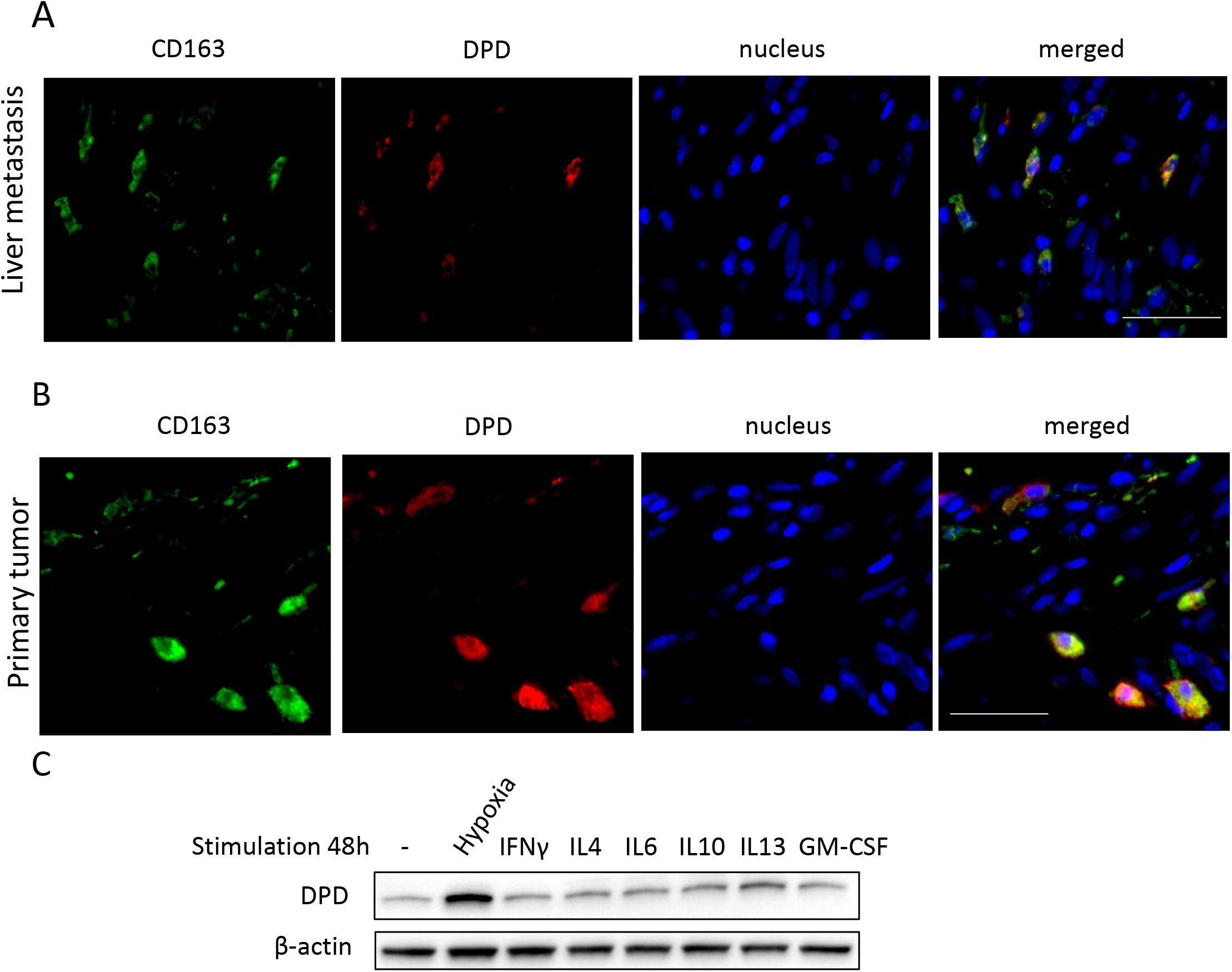
(A) Immunofluorescence staining in liver metastatic tissues. CD163 is in green, DPD in red, nuclei are stained by Hoescht in blue (n=4; scale bar= 50 μm). (B) Immunofluorescence staining in primary tumors. CD163 is in green, DPD in red, nuclei are stained by Hoescht in blue (n=4; scale bar= 50 μm). (C) Immunoblot analysis of DPD expression in human macrophages exposed to IL-4 (20 ng/mL), IL-13 (20 ng/mL), IL-10 (25 ng/mL), IFNγ (10 ng/mL), IL-6 (20 ng/mL) and GM-CSF (25ng/mL) compared to hypoxic condition for 24h (n=3). β-actin is used as control loading (n=3).

| Sex | Age | Comorbidities | synchronous<br>metastasis: 1 | Number of liver<br>metastasis | First line chemotherapy | Targeted<br>therapy | decease from<br>cancer | Desease free<br>survival<br>(months) | Total number of<br>treatment lines |
| --- | --- | --- | --- | --- | --- | --- | --- | --- | --- |
|  |  |  | metachronous<br>metastasis: 2 |  |  |  |  |  |  |
| F | 62 | / | 1 | 1 | Fluorouracil + Irinotecan | anti-VEGF | yes | 25 | 1 |
| F | 65 | diabetes, Hypertension | 2 | 2 | Fluorouracil + Irinotecan | none | yes | 73 | 3 |
| M | 61 | / | 1 | 2 | Fluorouracil + Oxaliplatin | anti-EGFR | yes | 63 | 3 |
| M | 64 | Cardiopathy, Hypertension, diabetes | 1 | 3 | Fluorouracil + Oxaliplatin | anti-VEGF | alive | 77 | 1 |
| M | 74 | Hypertension, asthma | 1 | 1 | Fluorouracil + Oxaliplatin | none | alive | 25 | 1 |
| M | 78 | / | 2 | 1 | Fluorouracil + Oxaliplatin | none | alive | 56 | 1 |
| M | 69 | / | 2 | 2 | Fluorouracil + Oxaliplatin | anti-VEGF | alive | 43 | 4 |
| M | 45 | alcoholism, hepatic cirrhosis | 1 | > 5 | Fluorouracil + Oxaliplatin | none | yes | 53 | 4 |
| M | 73 | Asthma | 2 | 1 | Fluorouracil + Oxaliplatin | none | alive | 142 | 1 |
| F | 48 | / | 1 | 2 | Fluorouracil + Oxaliplatin | anti-VEGF | yes | 66 | 5 |
| F | 45 | / | 1 | 2 | Fluorouracil + Oxaliplatin | none | yes | 35 | 2 |
| M | 57 | / | 1 | > 5 | Fluorouracil + Irinotecan + Oxaliplatin | none | alive | 42 | 1 |
| M | 72 | cardiac insufficiency, diabetes, chronic obstructive pneumopathy | 2 | 5 | Fluorouracil + Irinotecan + Oxaliplatin | anti-VEGF | yes | 76 | 5 |
| F | 76 | / | 2 | 1 | Fluorouracil | none | yes | 100 | 2 |
| M | 66 | Alcoholism | 1 | 2 | Fluorouracil + Oxaliplatin | none | yes | 37 | 2 |

